# Reversing Time Averaging and Reconstructing Extinction Rates with Approaches from Image Processing

**DOI:** 10.1101/408864

**Authors:** Niklas Hohmann

## Abstract

In this paper, the relation between the extinction rate and the rate of last fossil occurrences as well as the relation between the fossil occurrence rate and the time averaged fossil occurrence rate is examined. Both relations are described by the same mathematical operation. This operation is commonly used in image processing, where it generates a blurring effect. Therefore the rate of last fossil occurrences can be taken as a blurred version of the extinction rate, and the time averaged fossil occurrence rate as a blurred version of the fossil occurrence rate. This connection has different applications. It allows to study the patterns different types of time averaging generate or the patterns of last fossil occurrences generated by different extinction rates. More importantly, it opens the possibility to use algorithms from image processing that reverse blurring effects for geological applications. This can be used to reverse the effects of time averaging or to reconstruct extinction rates from the rate of last fossil occurrences.

## 1 Introduction

In this paper, the structural similarities between two fundamental biases on the fossil record are demonstrated and a possibility to reverse both using approaches from image processing is sketched.

The first of these biases is time averaging, causing fossils not to obey the law of superposition as it was formulated by Nicolas Steno in the 17th century, and therefore contradicting an axiom of stratigraphy. Due to time averaging, the age of fossils in the stratigraphic column is not only unordered, but signals based on fossil abundances get blurred, since fossils migrate into overlying or underlying stratigraphic layers and thereby average their fossil content[Kidwell et al., 1991]. The second bias is best explained by the Signor-Lipps effect[Signor and Lipps, 1982], which states that an extinction event will always appear more gradual as it actually was. This is since the knowledge of an extinction is not from the extinction of a taxon itself, but via the last fossil occurrence of this taxon. But the last fossil occurrence is only loosely connected to the actual extinction of the taxon, since it is heavily dependent on chance. Therefore the rate of last fossil occurrences *f_LFO_* is a blurred version of the actual extinction rate *f_ext_*, with the blurring generated by the loose connection of time of extinction and last fossil occurrence.

What unifies these two biases is that what can be observed, be it time averaged signal or rate of last fossil occurrences, is a blurred version of what is actually of interest (not time averaged data/extinction rates). If there is information available how this blurring is generated (the process that generates time averaging/the relation of last fossil occurrence to extinction), then the original signal can be reconstructed from the blurred signal. Reversing such blurring effects is a common problem in image processing, and image processing also offers a variety of solutions for this problem that can be used in this geological context.

One of the basic operations in image processing is a blurring or a smoothing effect. Many versions of these effects are based on a mathematical operation called a convolution, which combines a convolution kernel with the original picture to generate a blurred version of the picture. The convolution operation can be paraphrased as a continuous version of weighted moving averages, with the convolution kernel being the continuous equivalent of the weights. A common example for a convolution operation is the Gaussian blur, which uses a normal distribution as the convolution kernel (see fig. 2).

In many applications, such as microscopy, it is useful to assume that the recorded data is already blurred in a systematic way, which might for example depend on the type of microscope used. If information about this systematic blurring is available, algorithms can be applied to reconstruct the original, unblurred image. This procedure is called *deconvolution*, since it tries to reverse the blurring effects of a convolution. The result of the deconvolution is in general not unique (but see appendix B), and knowledge of the original blurring effect in the form of a convolution kernel is necessary.

It can be shown (see appendix A) that the mathematical relation of the time averaged rate of fossil occurrences *f_tavg_* to the rate of fossil occurrences *f_foss_* and the relation of the rate of last fossil occurrences *f_LFO_* to the extinction rate *f_ext_* is the same as the relation of a blurred image to its original version:

- The time averaged rate of fossil occurrences *f_tavg_* is a blurred (convoluted) version of the rate of fossil occurrences *f_foss_*
- The rate of last fossil occurrences *f_LFO_* is a blurred (convoluted) version of the extinction rate *f_ext_*

This connection is based on a duality in probability theory, where samples are taken as realizations of an abstract signal. The mathematical proof in appendix A shows that a model that is based on a relocation of samples on the sample level is reflected on the abstract signal level by a convolution of the signal (see fig. 1). Modeling time averaging and the Signor-Lipps effect as a relocation of the fossil occurrences and extinctions as done in section 2 and 4 is thereby reflected on the signal level as a convolution of the rate of fossil occurrences or extinction rate with a convolution kernel whose shape is determined by the type of relocation.

**Figure 1:**
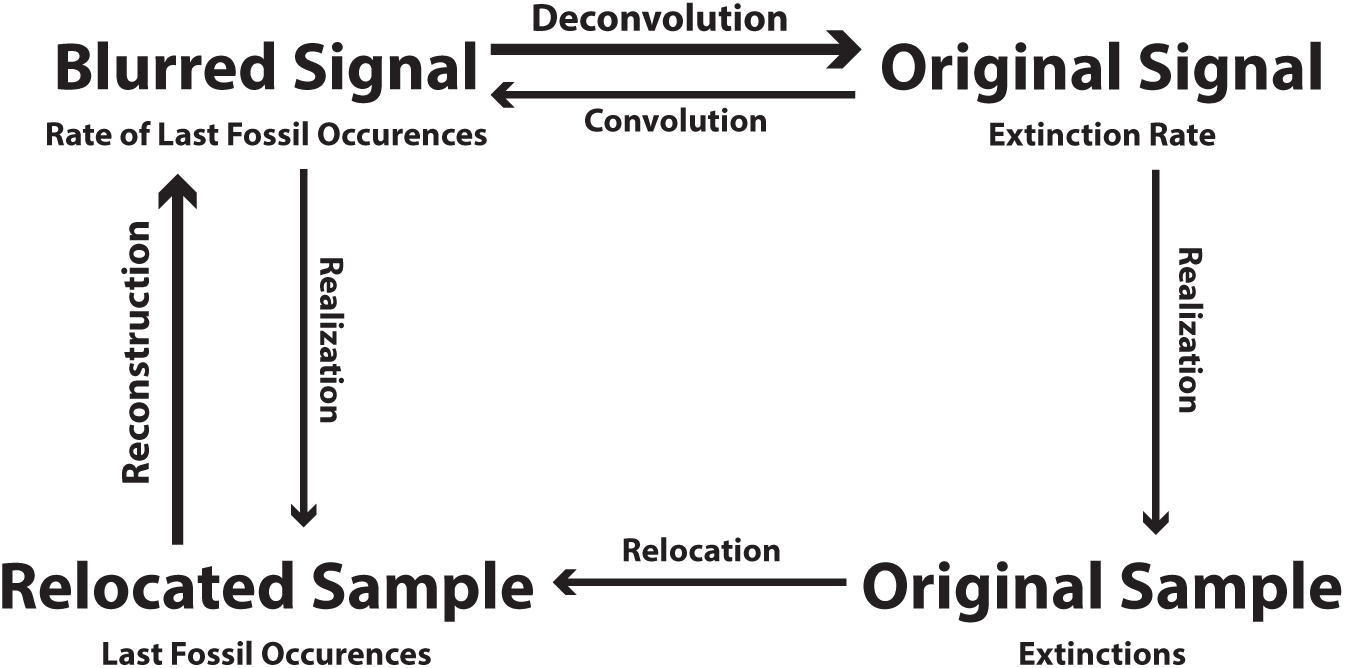
The overall structure proposed. The upper half is the signal level, the lower half the sample level. The connection between signal and sample is created by a realization of the signal, just as a random number is a realization of a random number generator. The model (M) describes how the original sample is transformed into the relocated sample. The proof in appendix A shows that the convolution is the equivalent description of this relocation on the signal level. Therefore it does not matter whether first the original signal is transformed into the blurred signal, which is then used to generate a relocated sample, or whether a sample is generated from the original signal, which is then relocated to generate the relocated signal.

**Figure 2:**
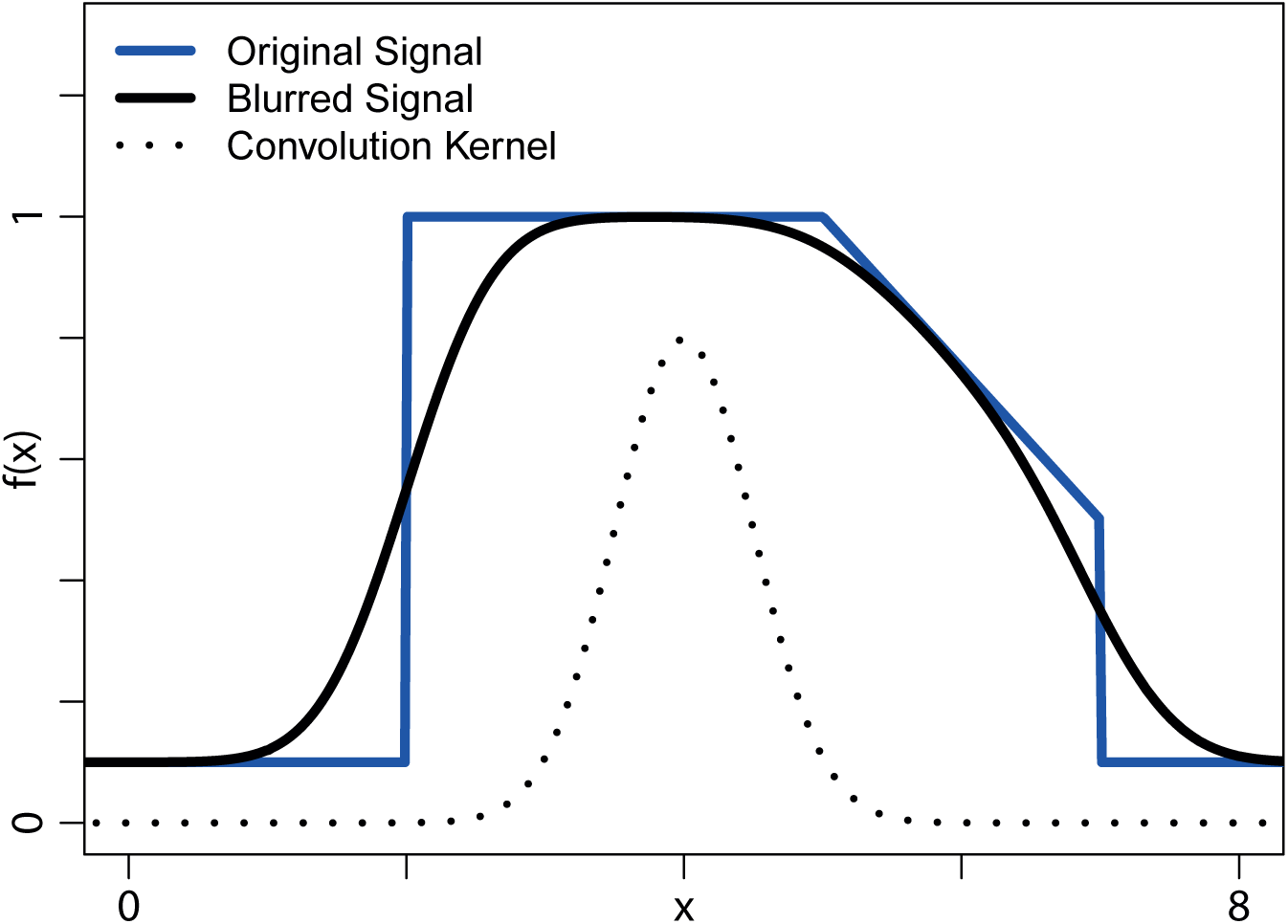
A discontinuous signal (dark blue), the convolution kernel (dotted), and the result of the convolution (black). Note that the sharp edges of the original signal are smoothed. The convolution kernel chosen here is the density function of a Normal distribution with standard deviation 0.5. The convolution displayed here corresponds to the Gaussian blur as it is implemented in most image processing programs.

So if knowledge about the convolution kernel that describes these blurring effects is available, the rate of fossil occurrences can be reconstructed from the time averaged rate of fossil occurrence and the extinction rate can be reconstructed from the rate of last fossil occurrences by applying a deconvolution algorithm.

## 2 Model Assumptions

*For clarity of language, only the case of time averaging is discussed here. The corresponding procedure and model assumptions for rates of last fossil occurrences and extinction rates can be found in section 4.*

In a first step, the fossils are abstracted as points on a line. This line can either represent stratigraphic height or time, dependent on the type of time averaging that is being modeled. The basic model can be described as

**(M)** Time averaging is a process of simultaneous destruction and relocation of fossils

This model is complemented by the following model assumptions:

**(TA 1)** The appearance of fossils is relatively rare, stochastically independent and the rate of fossil occurrences is given by a function *f_foss_*. This means that samples are realizations of an inhomogeneous Poisson point process (IPPP) with rate function *f_foss_*.

**(TA 2)** Destruction probabilities of fossils remain constant. Since the original number of fossils remains unknown, having a constant destruction probability is equal to no destruction at all. Therefore destruction of fossils will be neglected further on.

**(TA 3)** The relocation of fossils is random, but remains unchanged throughout time or with depth. This means that there is one probability distribution *P* that determines how much a fossil is being relocated relative to its original position.

These model assumptions are discussed in detail in section 6.1.

## 3 The Method

Assume that the following is given:

- The rate of time averaged fossil occurrences *f_tavg_*, as it can for example be reconstructed from time averaged data
- Knowledge of the distribution *P* that determines the relocation of the fossils.

Let *f_P_* be the density function (continuous case) or probability mass function (discrete case) of the distribution *P*.

Then *f_P_* is the convolution kernel that transforms the rate of fossil occurrences *f_foss_* into the rate of time averaged fossil occurrences *f_tavg_*:

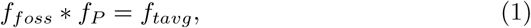

where * denotes the convolution. This equation is derived in appendix A.

The rate of fossil occurrences *f_foss_* can therefore be reconstructed as the deconvolution of the rate of time averaged fossils occurrences *f_tavg_* and *f_P_*:

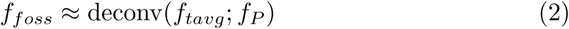

The overarching structure of these relations is displayed in fig. 1.

## 4 Extinction Rates and Last Fossil occurrences: Reversing the Signor-Lipps Effect

The case of extinction rates is identical to the case of time averaging, when last fossil occurrences are interpreted as random replacements of the actual extinctions. In this setting, the model assumptions from time averaging translate as follows:

**(EXT 1)** The extinctions follow an inhomogeneous Poisson point process with rate function *f_ext_*, and the fossil occurrences of each taxon follow an in-homogeneous Poisson point process

**(EXT 2)** Sampling effort is constant

**(EXT 3)** All taxa have the same rate of fossil occurrences *λ* close to their last fossil occurrence

These model assumptions are discussed in section 6.1. In the case of extinctions, the shape of the convolution kernel *f_P_* is alredy determined to be the density function of a exponential distribution with mean 1/*λ*, mirrored at the y-axis. This follows since the fossil occurrences are themselves Poisson point processes, which have exponentially distributed waiting times between fossil occurrences, and the memorylessness of the exponential distribution.

So just as in the case of time averaging, the rate of last fossil occurrences *f_LFO_* is given as the convolution of *f_P_* and the extinction rate *f_ext_*:

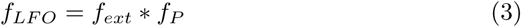

Therefore the extinction rate can be reconstructed as a deconvolution of *f_P_* and *f_LFO_*:

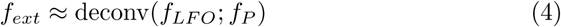

**Figure 3:**
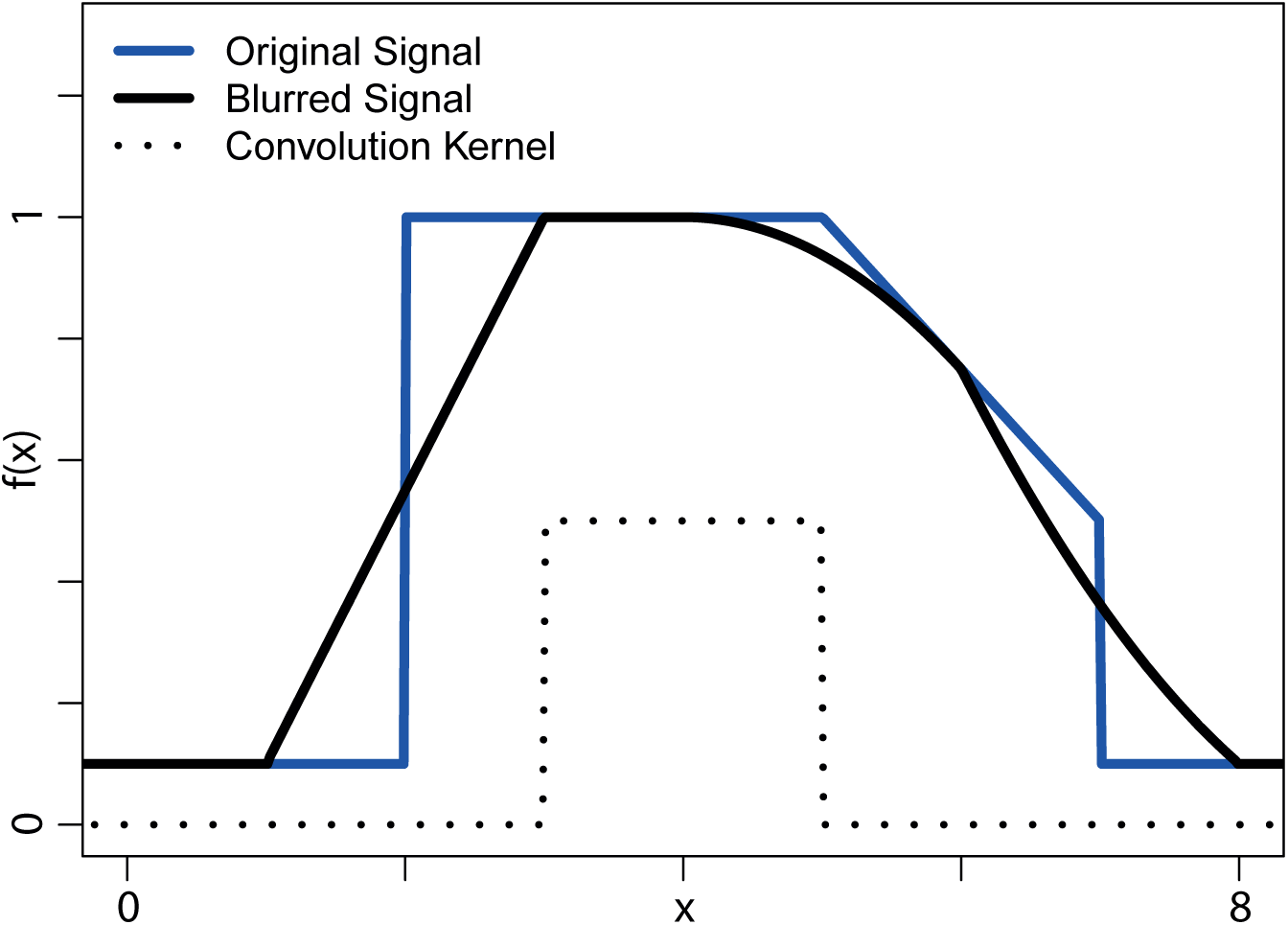
A discontinuous signal (dark blue), the convolution kernel (dotted), and the result of the convolution (black). The convolution kernel chosen is the density function of a uniform distribution with width 2. Although the original signal has been smoothed, some edges are still present in the blurred signal. This is since the convolution kernel is not as smooth as in fig. 2. This demonstrates that properties of the convolution kernel are reflected in the blurred signal.

**Figure 4:**
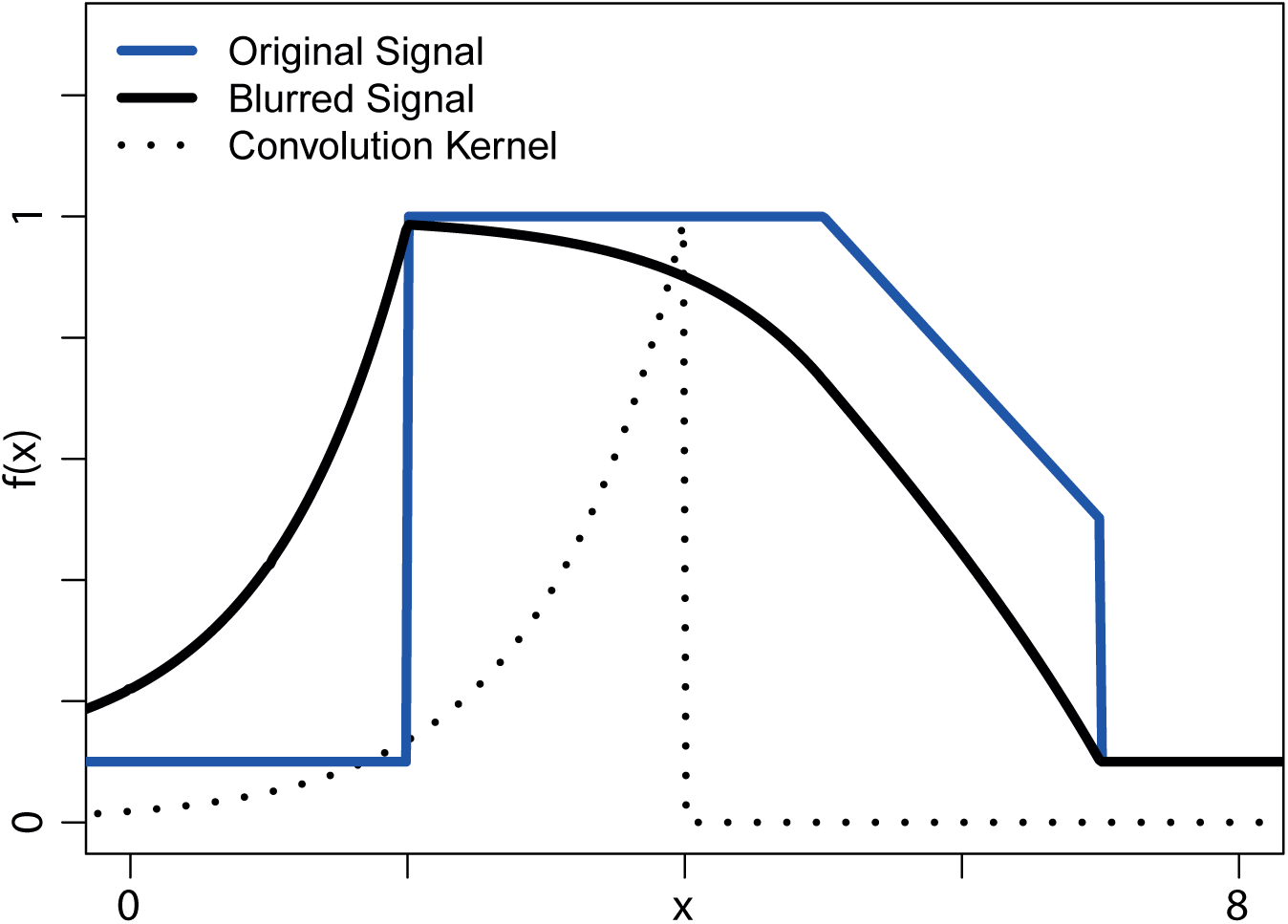
A discontinuous signal (dark blue), the convolution kernel (dotted), and the result of the convolution (black). The convolution kernel chosen here is the density function of an exponential distribution with mean 1, mirrored at the y-axis. The blurred function appears very asymmetrical. This is generated by the asymmetry of the convolution kernel. Note how similar very left of the blurred signal is to the convolution kernel. This similarity will in a geological context create the Signor-Lipps effect.

**Figure 5:**
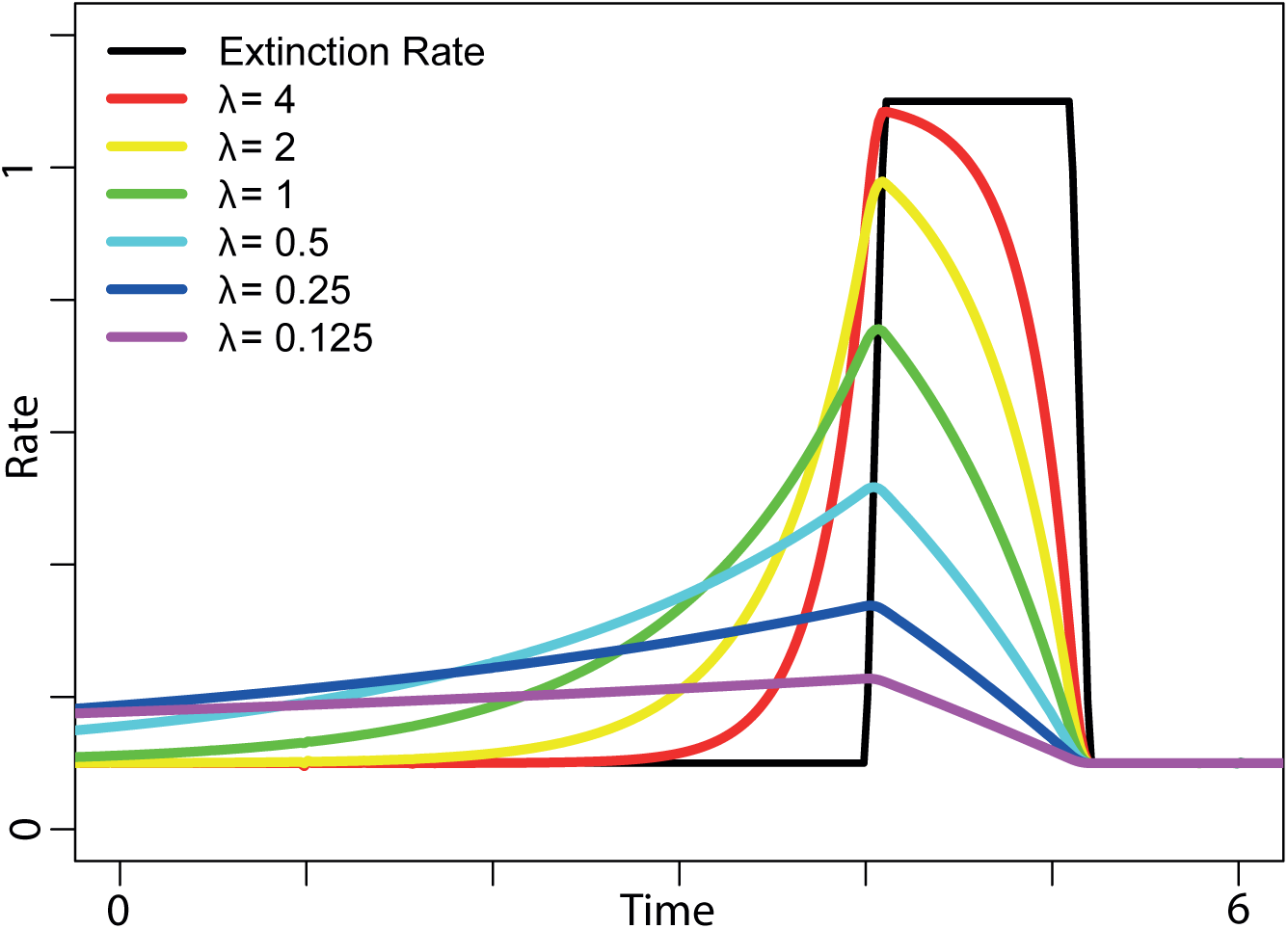
An extinction rate (black) and the derived rates of last fossil occurrences for different parameters *λ*. This parameter represents the rate of fossil occurrences of the taxa going extinct. As the rate of fossil occurrences decrease, the distance between last fossil occurrence and extinction increases up to a point where the rate of last fossil occurrences is so low that it does not resemble the extinction rate any more.

**Figure 6:**
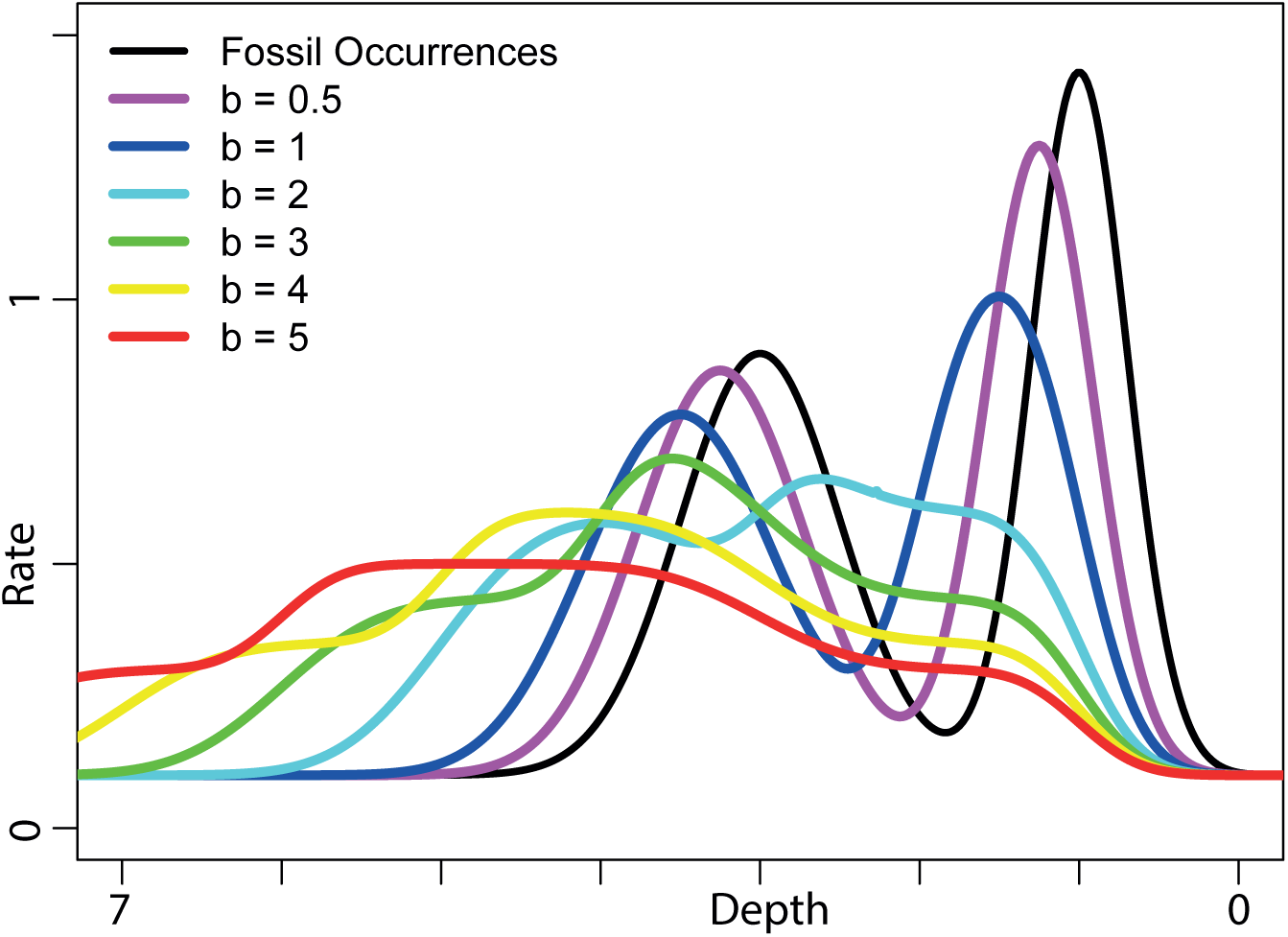
A rate of fossil occurrences (black) and different time averaged versions (in colors) of it. The convolution kernel is the density function of an uniform distribution from 0 to *b*. Note how the two distinct spikes in the original rate of fossil occurrences become more blurred until they cannot be distinguished anymore (*b* = 5, red)

The structure of the relations described here is displayed in fig. 1.

## 5 Examples and Figures

## 6 Discussion

The proposed model can not only be used to reconstruct extinction rates and reverse the effects of time averaging, but also to study how the effects of time averaging and the Signor-Lipps effect change under different extinction rates or different types of time-averaging and rates of fossil occurrences. For the case of only studying these effects, the model assumptions (TA 2)/(EXT 2) and (TA 3)/(EXT 3) are no longer binding if a brute force approach for the relocation based on the simulation of inhomogeneous Poisson point processes is used (see attached files of [Hohmann, 2017] for examples of code).

It is important to note everything presented here is based on the assumption that sequence stratigraphy is not interacting with any of the processes discussed. For applications, this is in general unrealistic. However the effects of changing deposition rates can be incorporated with the approach presented in [Hohmann, 2018]. This approach is based on the same mathematical concept and therefore compatible with the work presented here.

### 6.1 Model Assumptions

Assuming that all processes discussed here follow an inhomogeneous Poisson point process (TA 1 and EXT 1) is crucial for establishing the mathematical framework. This assumption is weak, since Poisson processes can be derived from an elementary sampling procedure that is based on the relative rarity and the independence of the events (fossils, extinctions, etc)[Meester, 2008, ch. 7]. Both relative rarity and independence are plausible assumptions for the paleontological processes discussed here.

#### 6.1.1 Model for Time Averaging

For time averaging, it is possible drop model assumption (TA 2) and to incorporate changing destruction rates of shells. For the sake of clarity this has not been done here. Whether model assumption (TA 3) holds is a priori not clear. If it does not hold, an exact reconstruction based on the mathematical framework developed here seems not possible. It might however be possible to establish error estimates for the reconstruction via deconvolution for cases where (TA 3) is violated.

#### 6.1.2 Model Assumptions for Extinctions

Weakening the assumption that sampling effort is constant (EXT 2) would require the development of a more advanced model, since changing sampling interacts with the model assumption (EXT 3). The assumption (EXT 3) about the constant rate is equivalent to the assumption (TA 3), since it makes sure that the relation between extinction and last fossil occurrence remains unchanged throughout time. It is not clear whether this condition holds, and checking it might be hard if fossil occurrences are rare. However it might be possible to establish an error estimate for the reconstruction via deconvolution for cases where (EXT 3) does not hold, just like in the case of time averaging.

### 6.2 The Perspective of Information Theory

In Information theory, the convolution is known to increase entropy [Chirikjian, 2009, p. 75] and to decrease information [Chirikjian, 2009, p. 78]. It is however not clear whether these properties hold in the cases discussed here. This is since the functions that are convoluted are not probability density functions, but representatives of a Poisson point process. They are therefore closer to very high-dimensional parameters than to probability distributions, and it is not clear whether the convolution of these parameters reflects any relevant information-theoretic operation.

But if these fundamental inequalities of information theory generalize into the context discussed here, this would mean that there is a general loss of information that can not be recovered, and reconstructions will to some point always be deficient.

Another approach from information theory that can be applied is looking at the relative entropy as described in [Hohmann, 2017] and use it to assess how the distinguishability of different signals changes through convolution. For a similar approach in the context of push forward measures, compare [Hohmann, 2018].

### 6.3 Remaining Questions

It remains to identify suitable deconvolution algorithms for the cases where the obvious approaches via Fourier transform or as sketched in appendix B fail. Since the nature of the convolution kernel is not a priori known in the case of time averaging, a second question is how to reconstruct these kernels for time averaging and how these kernels are linked to specific environments.

## 7 Acknowledgements

I would like to thank Adám Kocsis for his detailed feedback on the draft.

## Appendix A: The Connection Between Convolution and the Model

For the sake of simplicity, only the proof for the case of time averaging is sketched. The case for extinction rates follows mutatis mutandis, with *f_ext_* replacing *f_foss_*, *f_LFO_* replacing *f_tavg_*, and the model asumptions being replaced by their equally enumerated counterparts.

Model assumption (TA 1) states that the not time averaged sample is generated by an inhomogeneous Poisson point process *ξ* with intensity measure

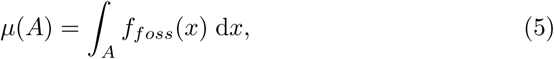

where *f_foss_* is the rate of fossil occurrences. So the number of fossils in an interval [*a, b*] follows a Poisson distribution with parameter *λ* = *μ*([*a, b*]), and the average number of fossil occurrences in the interval [*a, b*] is *μ*([*a, b*]).

Taking the basic model (M), the time averaged fossil occurrences are a *ν*-transform (in the sense of [Kallenberg, 2017, p. 73]) of the original process *ξ*, where *ν* is some Markov kernel.

With model assumption (TA 2), this Markov kernel contains no thinning component and therefore is a Markov kernel from ℝ to ℝ. Following theorem 3.2 in [Kallenberg, 2017], the *ν*-transform of *ξ* is a Cox process directed by *μ* · *ν*, where · denotes the product (sensu [Kallenberg, 2017, p. 33], but “composition” sensu [Klenke, 2008, p. 281]) of the measure *μ* and the Markov kernel *ν*.

Model assumption (TA 3) then states that *ν_t_* = *δ_t_* * *P*, where *δ_t_* is the Dirac measure on *t*. With Klenke [2008, lemma 14.27], it follows that

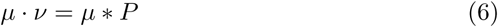

Therefore the Cox process describing the time averaged signal reduces to a Poisson process with intensity measure *μ* * *P*. This measure has by definition a density function of *f_foss_* * *f_P_*.

So if all model assumptions hold, the rate function *f_tavg_* of the time averaged process is given as the convolution of *f_foss_* and the probability density function *f_P_*:

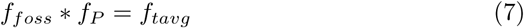

## Appendix B: A Unique Deconvolution

In the case of binned data, all functions reduce to series. These series can then be padded with infinitely many zeros on each sides and taken as elements of the sequence space *ℓ*^1^ (ℤ) with the corresponding norm ∥ · ∥_1_. This space is closed under the convolution and forms a Banach algebra with identity *e* = (…, 0, 0, 0, 1, 0, 0, 0, …), where the 1 is at the position with index zero. If *f_P_* is the convolution kernel, and

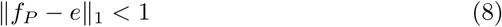

then the kernel *f_dec_* that describes the deconvolution is given as [Kaniuth, 2008, lemma 1.2.6]

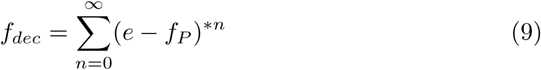

with the convention (*e* - *f_P_*)^*0^ := *e* and *x*^*n^ := *x*^*(*n*−1)^ * *x* being the n-fold convolution power. This means that

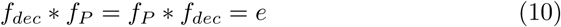

Since *f* * *e* = *e* for all *f* ∈ *ℓ*^1^ (ℤ), it follows that

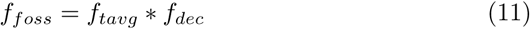

The result for the extinction rate follows mutatis mutandis.

Note that although the deconvolution kernel is uniquely determined in this case, this does not necessarily mean that *f_foss_* is positive and therefore a plausible reconstruction.

